# Data-driven microscopy allows for automated targeted acquisition of relevant data with higher fidelity

**DOI:** 10.1101/2022.05.09.491153

**Authors:** Oscar André, Johannes Kumra Ahnlide, Nils Norlin, Vinay Swaminathan, Pontus Nordenfelt

## Abstract

Light microscopy is a powerful single-cell technique that allows for quantitative spatial information at subcellular resolution. However, unlike flow cytometry and single-cell sequencing techniques, microscopy has issues achieving high-quality population-wide sample characterization while maintaining high resolution. Here, we present a general framework, data-driven microscopy (DDM), that uses population-wide cell characterization to enable data-driven high-fidelity imaging of relevant phenotypes. DDM combines data-independent and data-dependent steps to synergistically enhance data acquired using different imaging modalities. As proof-of-concept, we apply DDM with plugins for improved high-content screening and live adaptive microscopy. DDM also allows for easy correlative imaging in other systems with a plugin that uses the spatial relationship of the sample population for automated registration. We believe DDM will be a valuable approach for reducing human bias, increasing reproducibility, and placing singlecell characteristics in the context of the sample population when interpreting microscopy data, leading to an overall increase in data fidelity.

## Introduction

Over the last decade, advances in light microscopy, including faster and more precise motorized components, automated focusing and image acquisition, and more integrated computational power, have changed the field. Today, researchers are capable of collecting and analyzing thousands of images, accumulating millions of data points on cellular processes over a few hours. In parallel, we see an increasing focus on single-cell data, including at high spatiotemporal or super resolution level. Analysis and selection of such data could benefit from placing the cells in a population-wide context, with access to overall cell feature distribution, similar to how flow cytometry is routinely used. However, when studying cell populations in microscopy, researchers typically face a trade-off between cellular resolution and overall population assessment. It is thus technically challenging to put high-resolution single-cell data in a population-wide context.

Correlative imaging is one technique that aims at partially solving this trade-off. By correlating points between images captured in different modalities, the strengths of each technique can be combined. Such systems are typically tailored for the high-resolution to super-resolution domain, custom-built, and require specialized expertise. Because of the inherent spatial resolution of this domain, such systems are difficult to calibrate and force researchers into using single-system multi-modal imaging (1, 2), with on-stage fixation and labeling protocols (1,3,4). On the other hand, multi-system correlative imaging (5, 6, 7, 8, 9) is dependent on finding a transformation between the respective coordinate systems. While differences between the systems may prohibit off-the-shelf registration solutions (10, 11, 12, 13), the introduction of external cues such as markings (6) or calibration beads may influence the sample and obstruct image quality. Ideally, reference points should be derived from as little information as possible to achieve multi-system correlative imaging consistently.

At the other end of the spectrum, high content screening (HCS) has become a central method to assess large quantities of single-cell data in a population-wide context (14, 15, 16). With automated image analysis pipelines, HCS enables researchers to extract rich and unbiased information from datasets that would otherwise overwhelm any human operator (17, 18, 19, 20, 21, 22, 23). In other settings where high-spatiotemporal information is necessary and can be acquired, HCS has successfully been integrated (24, 25). The alternative to HCS in high-spatiotemporal settings, acquiring data from selected points, typically leads to the loss of population context and risk of bias, especially since data selection is often left to human operators. Integrating image analysis, including machine learning classification algorithms, into the data selection has reduced the overall bias of the acquired subsample. These solutions, referred to as feedback-microscopy or intelligent microscopy, allow for the high-throughput targeted imaging of cells of interest in high-spatiotemporal settings (26, 27, 28, 29). However, there is no framework capable of relating targeted high-resolution image data to the overall sample across multiple modalities and platforms.

Here, we present a general microscopy framework, data-driven microscopy (DDM), which uses population-wide data to improve and control microscopy and enables cross-experiment image data validation. A data-independent acquisition phase performs high-throughput imaging and generates a population-wide phenotype assessment. This data includes the relative coordinates of each data point for the system, which feeds into the automated data-dependent acquisition of selected phenotypes. Combining the two orthogonal data sets yields population-wide data with high-fidelity object characterization data. DDM also inherently includes what constitutes an objective representation of the sample population. The general DDM framework applies to any motorized microscope, and all steps can be fully automated, including phenotype targeting. As a proof-of-principle, we apply DDM to target and enhance HCS quantification of multi-labeled cells, carefully assess cancer cell migration phenotypes using live feedback microscopy, and perform automated correlative microscopy across different microscopy systems, in all cases without human input with regards to data acquisition. Placing the combined data in a population context yields a more robust, reproducible, and efficient method for selecting, acquiring, and correlating data in microscopy.

## Results

### Data-driven microscopy as an approach for automated targeted image acquisition of relevant data

We first asked what a representative cell is in a given sample. To address this question, we asked an experienced microscopist to acquire images of fixed triple-transfected cells in a usual manner (Fig. 1a, i). Next, we acquired a scan of the whole sample and quantified morphological features of the different channels in all cells (N=169988) (Fig. 1a, ii). Feature analysis demonstrated an expected feature heterogeneity across the sample (Fig. 1a, iv), and the cell heterogeneity is visualized as a UMAP of the combined features (Fig. 1a, v). The population analysis allowed us to place the high magnification – and manually targeted – cells in a feature distribution and population-wide context. Mapping the selected cell data onto the population data revealed an unconscious bias in the manual acquisition that did not represent the whole population of cells in the feature distribution (Fig. 1a, iii, iv) and UMAP space (Fig. 1a, v, vi). This experiment highlighted common issues with how light microscopy imaging is performed today, which also opened an obvious opportunity. We hypothesized that it should be possible to reverse the order of the approach and use population-wide data to target either truly representative cells or interesting sub-populations.

**Fig. 1.**
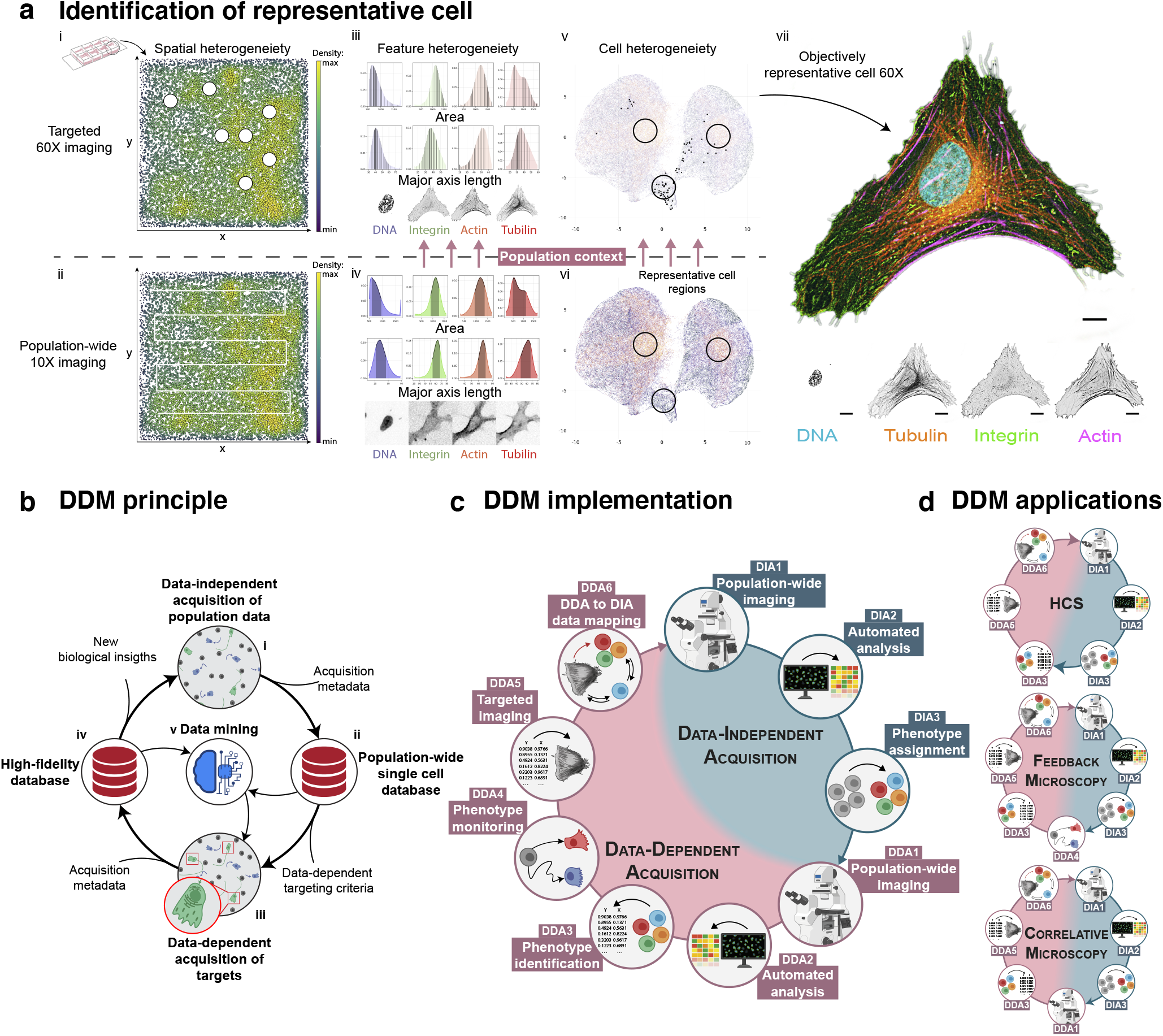
Data-driven microscopy as an approach for automated targeted image acquisition of relevant data. **a.** Objectively representative cells can be identified through the combined use of data from two traditional image acquisition strategies. HeLa cells were labeled with fluorescent tubulin, integrin, actin, and DNA. Targeted imaging (i) typically involves the manual selection of cells of interest. This leads to selected, high-resolution data (N=78 cells) that lack population context (ii). Population-wide automated imaging leads to a complete dataset (iv; N=169988 cells) of all present phenotypes but lacks high-resolution information about cells of interest (iii). There exist several types of heterogeneity in the data (UMAP space of multidimensional collapsed cell features; v). Combining the two orthogonal data sets provides the population context (v, vi) to identify features that would describe a representative cell (vii). **b.** Principle schematic of data-driven microscopy (DDM). A data-independent acquisition step (DIA; i) acquires population-wide data on a single-cell level. This data is stored in a database for sample health validation and future reference (ii). A data-dependent acquisition step (DDA; iii) uses the data stored in the database to target phenotypes of interest for high-fidelity, high-intelligence smart microscopy imaging. High-fidelity data captured in a population-wide context can lead to new biological insights which drive future experiments (iv). The combined high-fidelity data and population-wide single cell data has the potential for data-mining and machine learning training algorithms that can drive future DDA experiments (v). **c.** A schematic representation of DDM implementation. DDM consists of two imaging strategies, with indicated modules that when combined lead to both high-resolution and population-wide data. Through DIA, whole-population, single-cell imaging is coupled with automated image and data analysis pipelines. This results in a seamless and unbiased data stream that the operator can explore. DDA can perform targeted acquisition at higher fidelity by characterizing cells of interest in the data. Through population-wide, low magnification imaging, phenotypes are monitored, and if found, the system performs an automated switch to high-fidelity imaging (e.g., high magnification, high sensitivity, or multiparameter imaging). **d.** Examples of different applications suitable for data-driven microscopy (DDM) and what modules of DDM would be used to run the various pipelines.

The principle of DDM is based on two general imaging strategies linked by a server-based database to achieve automated population-wide characterization or targeted single-cell high-magnification imaging (Fig. 1b). The implementation of DDM can be seen as sequence of modules (Fig. 1c). The first module is data-independent acquisition (DIA), where whole populations are imaged in low magnification and analyzed at a single-cell level. From this low-resolution image data, phenotypes are typically characterized by morphology, signal, or event-based interactions but can be any feature that can be extracted from the image data.

The second strategy, data-dependent acquisition (DDA), aims at the targeted acquisition of specific phenotypes. Initially, population-wide single-cell data is acquired and assessed in real-time using an integrated microscope-server solution (Supplementary Fig. 1-2). This data is normalized according to previously acquired data through DIA and DDA. If present, the positions of phenotypes of interest are recorded until the criterium for triggered high-magnification imaging is met. This criterium can either be a satisfied count of observed phenotypes, cellular events, or a combination. By obtaining low-and high-magnification single-cell data and low-magnification population wide-data, high spatiotemporal characteristics seen in the phenotypes of interest can be inferred onto phenotypes not yet imaged in high magnification. Taken together, a synergistic relationship can be established between DIA and DDA, increasing the fidelity of the data collected through each strategy. As a proof-of-concept, we apply DDM to create data-driven enhanced applications of HCS, live feedback microscopy, and correlative microscopy (Fig. 1d).

### DDM enables targeted high magnification imaging of multi-labeled subpopulations based on data-driven criteria

To test whether DDM could improve HCS, we developed a data-driven enhanced variant of HCS (Fig. 2a) whereby automated sub-sampled high-magnification data could curate the low-magnification population-wide data (Fig. 2b). The goal was to assess transfection efficiency by targeting cells transfected with multiple plasmids. We acquired the entire cell populations in 16 wells (Fig. 2c) and characterized the signal of each plasmid at a single-cell level. Each cell was categorized (non-, single-, double-or triple-transfected) according to a signal threshold per channel. Figure 2d shows an example of a field of view and cells from the respective automatically assigned transfection categories. The normalized intensity in each channel across the whole population of cells is shown in Figure 2e. The distribution of cells (N=109065) with the low-resolution data was found to be heavily skewed towards non-transfected (41.9% ± 1.0%) or single-transfected (53.5% ± 1.0%). Only small fractions of cells were identifiable as expressing two plasmids (4.6% ± 0.1%) or more (<1%) (Fig. 2f).

**Fig. 2.**
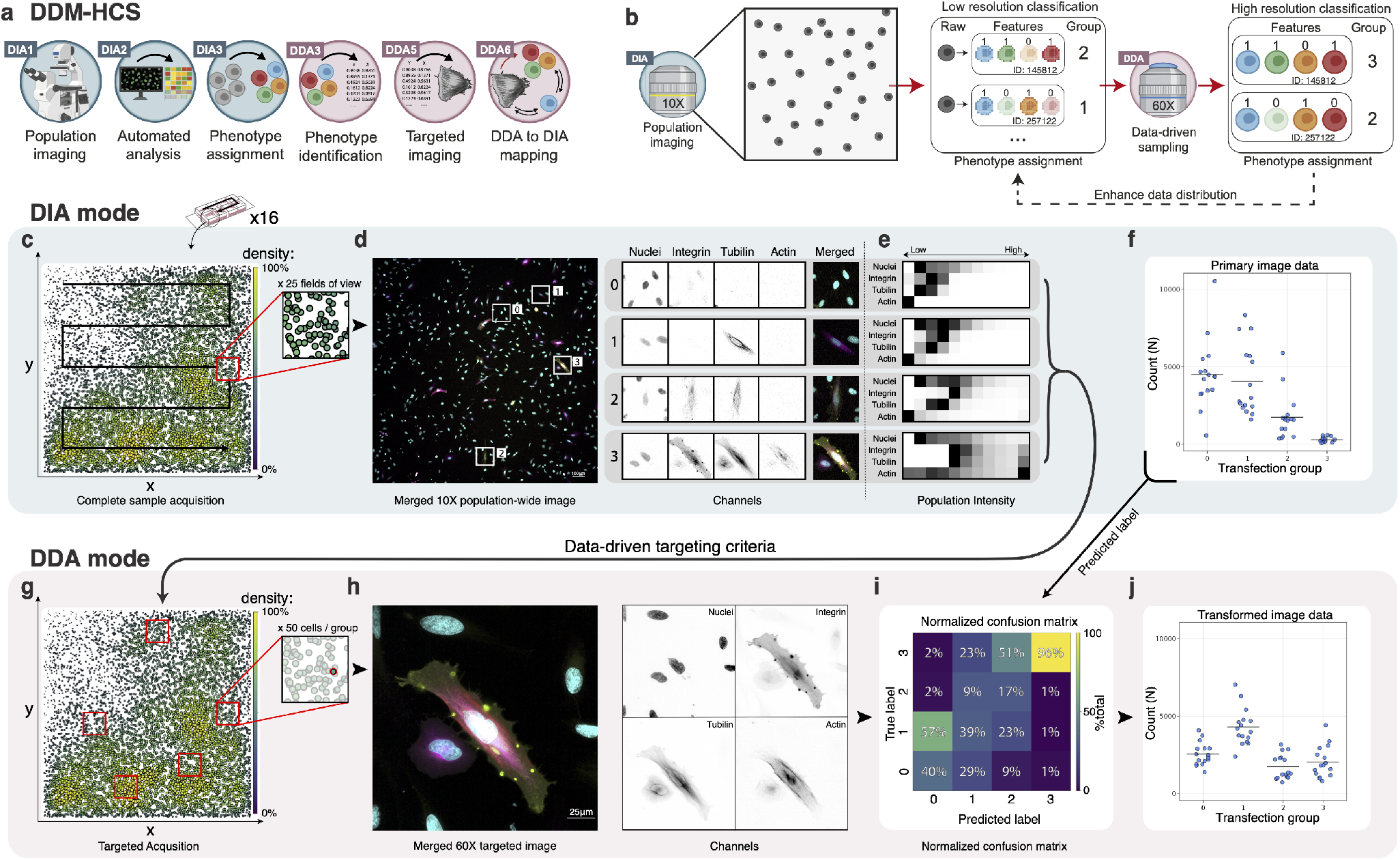
DDM enables high magnification imaging of targeted multi-labeled subpopulations. **a.** Illustration of the DDM setup used when studying multi-metric distributions of cells. **b.** Illustration of how DDM can increase data fidelity through the combination of DIA and subsampling of cells for DDA. **c.** The distribution of transfected HeLa cells in a sample imaged through DIA in 10X. **d.** Examples of negative, single, double, and triple transfected cells. **e.** The intensity of each channel per category normalized on the population-wide data and binned into quantiles. The rightmost (11th) pixel represents outlier cells (> 99th quantile). See also Supplementary Fig. 3. **f.** Quantification of transfection distribution of cells (N=109065) from different wells in each transfection category. **g.** In DDA, 50 cells per category were sampled for targeted high-magnification imaging. **h.** An example of a DDA-targeted triple transfected cell in 60X magnification. **i.** Confusion matrix of manually verified cells and the classification by the DIA algorithm (N=278). The heatmap represents the accuracy of the classifier. **j.** Confusion matrix-corrected DIA data of transfected cells (N=109065).

Following the DIA procedure, an even distribution of cells (N=50) from each category was automatically targeted for imaging in DDA (Fig. 2g). Figure 2h shows an example of a targeted triple-transfected cell in high-resolution (60X magnification); in this case, the same cell (nr 4) as shown in Figure 2d. The resulting high-resolution data from DDA was used to test the accuracy of the targeting criteria for each group. The resulting confusion matrix of predicted and manually curated cells (N=278) revealed an overly conservative prediction by the categorization during DIA (Fig. 2i). The prediction of single-transfected cells in DIA was inaccurate, as non-transfected cells were confirmed to be single-transfected 57% of the time compared with the predicted 29%. The DIA analysis module also misclassified triple transfected as double transfected (51%) or single transfected (23%). Triple transfected cells were accurately predicted (96%). With the confusion matrix, the inaccuracy in classification during DIA could be accounted for, and the distribution of categorized cells could be corrected (Fig. 2j). The transfection data illustrates that low magnification DIA (like HCS) is useful for initial acquisition and targeting of population data, but with clear misclassification of image data. However, after the DDM-enhanced analysis with high-resolution DDA data, the original DIA data could be corrected, increasing data fidelity, resulting in a large (N=109065), high-quality data set. Thus, we conclude that DDM can accurately characterize the distribution of multi-labeled cells through DIA, like traditional HCS, and subsequently perform high-magnification targeted imaging of subpopulations through DDA, leading to higher fidelity HCS through data-driven enhancement.

### DDM allows for high-spatiotemporal adaptive feedback microscopy of migratory subpopulations

To test the capacity of DDM on live feedback microscopy, we decided to study cancer cell migration at high spatiotemporal resolution. An adaptive feedback microscopy module was developed and implemented with DDM (Fig. 3a-b). Through DIA, H1299 mKate-paxillin cells (N=24940) were imaged every 10 minutes for 6 hours, and single-cell migration was characterized. The parameter space was collapsed using UMAP analysis to categorize the migration modes. To investigate this space, single-cell migration tracks from different regions were overlayed onto the UMAP (Supplementary Fig. 4). As expected, a wide range of phenotypes in terms of migratory behavior were present. To investigate further, we grouped the cells into their top and bottom percentiles (>90th and <10th, respectively) in terms of mean migration speed. This separation revealed distinct phenotypes, with the bottom percentile containing different migratory behaviors compared to the longer and generally smooth tracks in the top percentile (Fig. 3c). Grouping the cells based on their meandering index also showed a distinction between the respective groups. In the bottom percentile, cells were migrating around their original starting position, some of them in long smooth tracks. The top percentile migrates greater distances than the bottom percentile and in straighter paths (Fig. 3d). So far, analysis of the low-resolution DIA data shows a wide heterogeneity in the migratory behavior of the cells in the population.

**Fig. 3.**
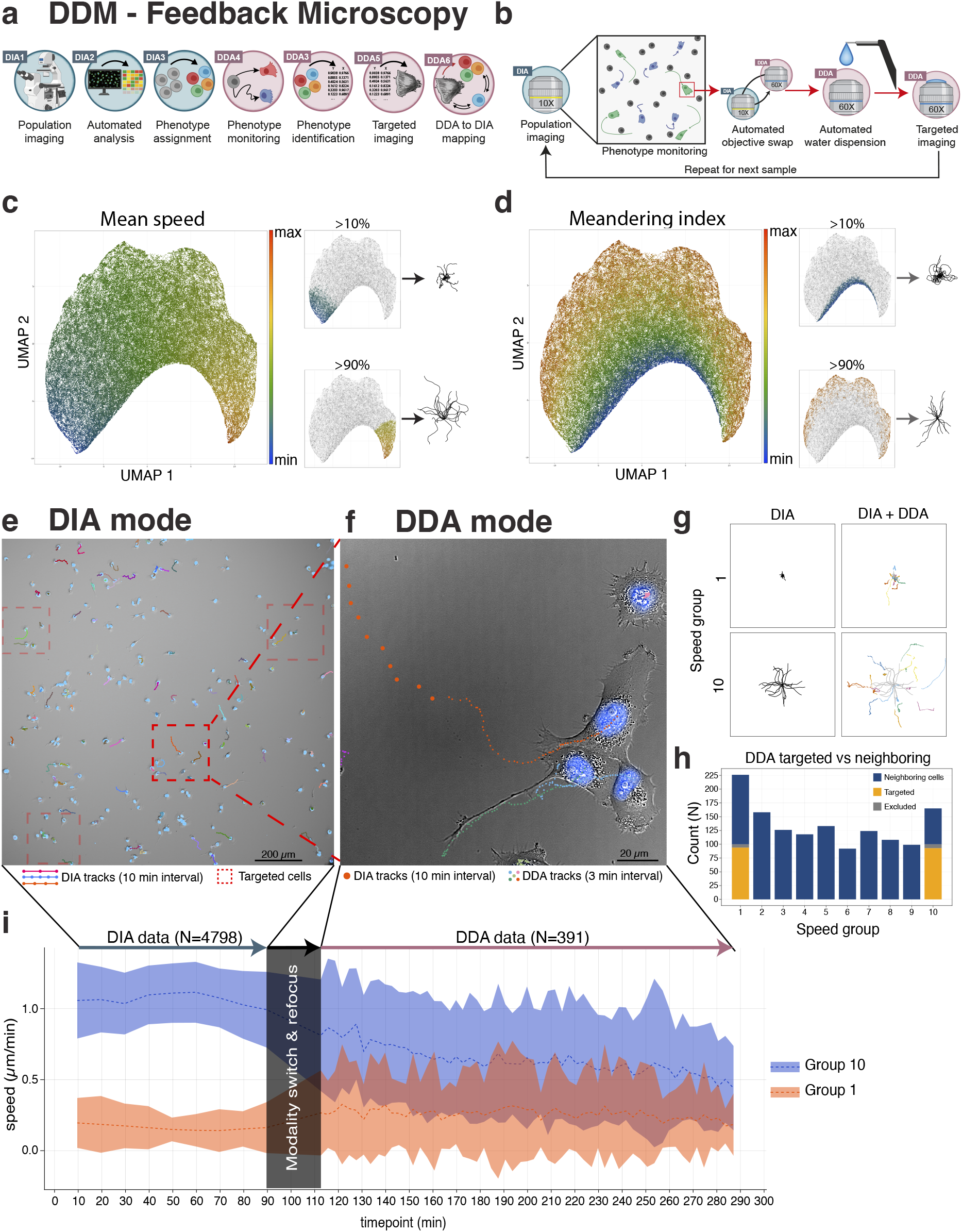
DDM allows for the characterization and high-spatiotemporal imaging of migratory subpopulations. **a.** Illustration of the DDM setup used for live-feedback microscopy, **b.** In DIA, cells were imaged every 10 min and analyzed post-acquisition. Cells were analyzed in multiple migration properties and categorized into slow and fast (<10th and >90th quantile respectively) in terms of mean speed. In DDA, subsampling of the two categories for high magnification imaging was aided by an automated water-dispenser. **c**,**d** UMAP space colored using mean-speed (**c**) and meandering index (**b**) revealed distinct migratory phenotypes. **e**,**f** Example image of cells migrating in different speeds in DIA (**e**) and a targeted cell in DDA (**f**) migrating in the top percentile with overlayed coordinates from the DIA and DDA (large and small markers respectively). Highlighted are fast (group 10; top left, top right, and middle rectangle) and slow (group 1; bottom left rectangle) cells in terms of mean speed. **g.** Cell migration was continuously tracked between DIA and DDA. **h.** From the DDA datasets, neighboring cells (blue) to the targeted cells (yellow) significantly increased the overall cell count. A small portion of the cells (gray) were excluded during analysis due to autofocus or tracking error. **i.** Speed variation over time during DIA (N=4798) vs DDA (N=391) for the two speed groups.

For understanding cell migratory behavior in more detail, we decided to target cells for data-driven imaging at high magnification. Through DIA, cells were imaged every 10 min for 100 min total, and single-cell migration was characterized. Figure 3e shows cells successfully tracked for the duration of the DIA. Since mean-speed as a metric previously displayed distinct migratory phenotypes in the extremes (Fig. 3c), we decided to sample (N=100) the fastest and slowest (>90th, <10th percentile, respectively) migrating cells for DDA. Figure 3f shows a targeted fast migrating cell, in this case, the same cell as one of the highlighted cells in Figure 3e. In DDA, images were taken every 3 min for 3 hours and the speed was monitored. Because both DIA and DDA were performed on the same system, the continuous tracking of cells between modalities could easily be performed (Fig. 3g). This could also be done on neighboring cells next to the targeted cells, which increased the overall dataset of fast and slow migrating cells to 391 (165 and 226, respectively, Fig. 3h). The final dataset also included 958 unique observations of cells in the intermediate groups (10th-90th percentiles), which further increased the total sample size to 1336. The increased spatial and temporal resolution of the DDA data resulted in a much more detailed tracking of cell migration, clearly seen in the increased variance of both groups in DDA (Fig 3i). In addition, although we see a decrease in the mean speed of the fast-migrating cells in DDA, they were continuously faster than the slow-migrating cells, indicating that most of the cells in each targeted group maintain their speed profile over the duration of the experiment. Taken together, DDM enables monitoring cells in a population-wide context in combination with an automated targeted high-spatiotemporal acquisition, resulting in increased overall fidelity and integrity of the data.

### DDM allows for multimodal correlative microscopy using cell-based coordinate transformation

DDM inherently collects information about the sample population, including the coordinates of each object in the population. We realized that this information can be used to infer the relative position of all cells in a sample, and thus theoretically allow for image registration across microscopy systems. Since the relative position of cells is independent of the imaging system, all that is needed for identifying cells across systems is to acquire a few fields of view in the new system and then calculate the coordinate transformation needed to relate the cell positions (Supplementary Fig. 6). This data-driven approach to correlative microscopy obviates the need for specialized markers and cumbersome image registration routines and can compensate for misplacement of the sample in the microscope, as the samples do not need to be in the same place for the coordinate transformation to work.

As a proof-of-concept for data-driven based correlative microscopy, we aimed to expand upon the experiment in Figure 3 with correlative SIM and TIRF microscopy and explore the focal adhesion (FA) properties of migratory H1299 subpopulations in more detail (Fig. 4a). Wildtype H1299 cells were imaged for 6 hours every 10 min and characterized on a wide-field microscope using DIA. We fixed and permeabilized the cells during the last time point, and subsequently stained the cells for high-resolution imaging (Fig. 4b). On two secondary systems (SIM and TIRF microscopes in separate buildings), spatial relationships between objects were recorded, and the DIA data was converted and mapped onto the coordinate system of the corresponding microscope (Supplementary Fig. 6). The correlation of sample coordinates between microscopes was achieved without manual adjustment and simply involved placing the sample in the new system and acquiring correlative data of data-driven targeted cells. The slowest (<10th percentile – mean speed: 5.64 ± 1.26 μm/h) and fastest (>90th percentile – mean speed: 43.20 ±8.04 μm/h) populations were sampled, and we targeted 50 cells per group for high-magnification imaging. Figure 4c exemplifies the same cells imaged on separate wide-field, SIM, and TIRF microscopes.

**Fig. 4.**
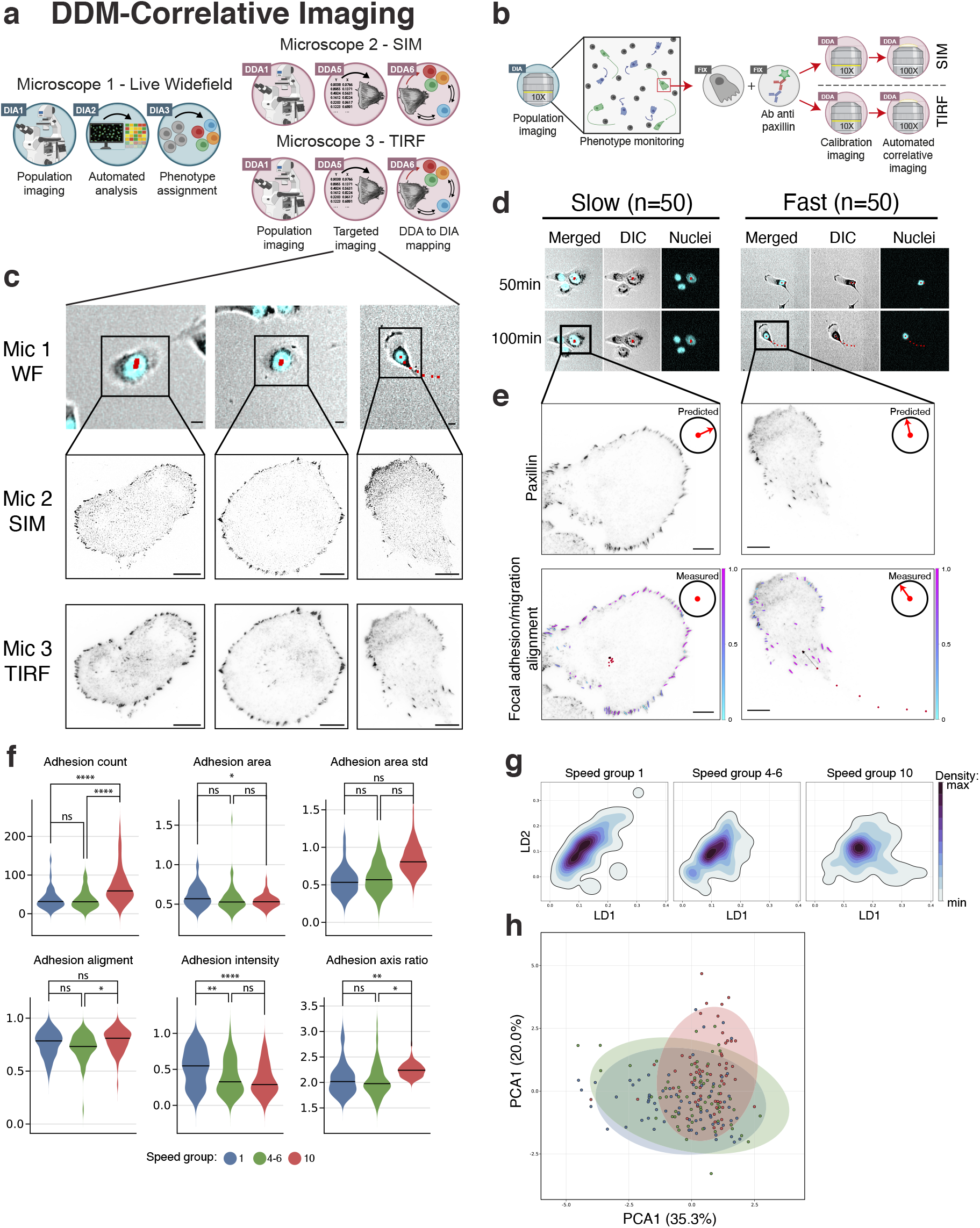
DDM allows for targeted correlative imaging of migratory phenotypes. **a.** Illustration of the DDM setup used for correlative imaging. **b.** Migration of H1299 cells was tracked through DIA. Following DIA, an intermediate fix and antibody labeling step was performed, and the dataset was calibrated for correlative imaging in SIM and TIRF microscopy (Supplementary Fig. 6). **c.** Example images of cells in three different microscopy systems (widefield live imaging, DIC + Nuclei –top row; SIM microscopy, Paxillin – middle row; TIRF microscopy, Paxillin-bottom row). **d.** Example images of fast and slow (>90th and <10th percentile, respectively) migrating cells in terms of mean speed acquired through DIA (N=50 per group). **e.** Correlative imaging of paxillin in the targeted cells. DIA data allows for the alignment of the focal adhesion (FA) major axis to the migration vector prefixation (bottom row, N=134). **f.** Violin plots of different properties for speed group 1 (<10th percentile, N=74, blue marker), group 4-6 (40th-60th percentile, N=65, green marker) group 10 (>90th percentile, N=60, orange marker). **g**, **h** separation of speed groups in linear discriminant analysis (**g**) and principal component analysis (**h**). Scale bars are 10 μm. Statistical significance was assessed using two-way ANOVA with Tukey multiple comparisons test. * denotes p ≤ 0.05, ** for p < 0.01, *** for p < 0.001, **** for p < 0.0001. ns not significant.

Time-resolved information, such as the direction of cellular migration, is unknown when performing fixed, high-resolution imaging. The overall shape of a cell can be used to predict the direction of migration when the sample is fixed, but this type of classification is highly uncertain. By combining migration data from DIA (Fig. 4d) with high-resolution data of FAs from DDA, we could correlate migration data from live imaging with high-resolution FA properties (Fig. 4e). The alignment of the major axis of the FAs to that of the direction of the cell movement in the last 10 min of migration was used to calculate the orientation. A comparison to shape-predicted migration vectors demonstrates the strength of having access to the true migration data, where the slow-moving cell would have been inaccurately classified otherwise.

Combining data from different microscopes allows for cross-correlative analysis, potentially revealing relationships that cannot be seen when only having access to data from one system or approach. In Figure 4f we grouped the FAs based on the respective speed group of each cell. We saw a significant increase in the number of FAs per cell, as well as a strong decrease in the adhesion localized paxillin signal intensity in fast migrating cells (group 10). We also see a small increase in the size of FAs (area) and a change in their shape (elongated axis ratio) of slow migrating cells (group 1). Multivariate statistical analyses of FA and cell properties showed a shift between the groups in linear discriminant analysis (Fig. 4g) and principal component analysis (Fig. 4h). This analysis shows that, while they share many characteristics, migratory subpopulations can be identified and tracked by DDM, and that they maintain certain properties over time. Overall, this automated application of correlating live-cell data to super-resolution imaging in multiple systems highlights the power of DDM and the benefit of using a data-driven approach to microscopy.

## Discussion

The operator dependency within microscopy leads to unintentional bias in data, difficulties in reproducing results, and potential misrepresentation of biological relevance according to the whole sample population. We have developed a new methodology for image data acquisition and selection, DDM, which uses a combination of automated multimodality imaging. Through DIA, we acquire population-wide data and profile single-cell phenotypes in the context of the population distribution. Using this data, we can acquire further data with other imaging modalities (DDA) that provide additional information about targeted phenotypes of interest. DDM is synergistic in that DDA complements DIA data, making both more relevant, leading to enhanced fidelity of the combined image data.

As proof-of-principle, we have used DDM with Nikon hardware to identify and profile representative cells in multilabeled samples and perform advanced characterization and multimodal correlative acquisition of cell migration subpopulations. The framework is compatible with any microscope as long as the controlling software can send images to the server or invoke external programs to achieve this. Upon identifying phenotypes of interest, we have automatically acquired additional images of these targets and assessed their relevance according to the population-wide data. We believe that these experiments have highlighted the versatility and benefits of DDM compared to traditional microscopy approaches.

DDM increases the throughput of biologically relevant data. The novelty of DDM compared to traditional HCS methods (15, 17, 30, 31) is that the image acquisition can easily be adapted according to the questions being asked and based on the population characteristics. The improvement is especially true for high-resolution applications and is exemplified through the transfection experiment we performed. The question was trivial, but the results were clear. If a traditional HCS approach had been used, it would have resulted in a large low-quality data set with misrepresented transfection distribution. If a traditional high-magnification approach had been used, it would have resulted in a small sampling of high-quality data with an uncertain transfection distribution. DDM uses the best aspect of each modality and controls acquisition in an automated and efficient manner, resulting in large data sets with high fidelity.

DDM inherently provides information on what constitutes representative objects in the sample population. Cells can be considered representative when they are placed in the context of the population feature distribution; it also opens the possibility to do analysis and targeting of outlier cells, as we did with the cell migration analysis, where we studied the slowest and fastest migrating cells in more detail. Since DDM essentially provides a population fingerprint for each experiment, it makes it less prone to human error and bias and could provide a basis for automated curation of large datasets (32), making it a powerful approach for AI-based analysis.

One of the most evident benefits of DDM comes when correlative microscopy is performed across different microscope setups. We easily acquired the correlative data sets with DDM, and the same cells were identified and imaged across wide-field, SIM, and TIRF microscopes placed in different labs. This allowed us to combine time-resolved single-cell data with specialized high-resolution spatial data and then cross-correlate the data. This correlative aspect alone opens for studies of many questions that are very difficult or require specialized equipment to address. An obvious future application of DDM would be different spatial omics solutions, where samples can be removed from the stage and easily automatically registered after offline processing.

To increase introspection and reproducibility, DDM inherently logs all operations performed, as well as the state of the running experiment, providing clear status updates to the user. The ability to fully interoperate the popular programming language Python allows DDM to be easily integrated into other already existing workflows and methods. More advanced image analysis, such as machine-learning approaches (33, 34, 35, 36), should make it possible to map the data collected through DDA backward onto the DIA data. In the future, this could potentially remove the need for DDA for specific applications. A similar development has been shown within the mass spectrometry field (37), where DIA data can be mined after the fact to provide high-quality data for new questions. In summary, we believe that DDM offers a useful framework for a more robust and unbiased acquisition of high-fidelity microscopy data.

## Methods

### Cell culture

The cell lines herein are not members of the ATCC list of commonly misidentified cell lines. All cells were maintained and used between passages 5-25. Human cervix epithelioid carcinoma cells (HeLa; Sigma Aldrich) were cultured in Dulbecco’s Modified Eagle’s (DMEM; Gibco, Thermo Fisher Scientific) supplemented with 10 % heat-inactivated fetal bovine serum (FBS) (Gibco) at 37 °C in a humidified 5 % CO2 incubator. For imaging, μ-slide 8-well glass-bottom slides (Ibidi) were coated with 2 μg/mL fibronectin human plasma (FN) (Sigma Aldrich) and incubated for 30 min at 37 °C. Cells were transfected in suspension ( 70 000 cells/mL) with 500 ng/plasmid using Lipofectamine 3000 (Invitrogen) according to the manufacturer’s protocol. All plasmids used are listed in Table 1. Cells were plated at 20 000 cells cm2 in each well and incubated for 48 hours before fixing with 4 % paraformaldehyde (PFA; Thermo Scientific) for 15 min at room temperature (RT).

H1299 cells expressing mKate-Paxillin and H1299 WT were kindly provided by Staffan Strömblad. H1299 cells (mKate-Paxillin and WT) were cultured in Roswell Park Memorial Institute (RPMI 1640; Gibco) supplemented with 10 % heat-inactivated FBS at 37 °C in a humidified 5 % CO2 incubator. For imaging, μ-Slide 4-Well Ph+ Glass Bottom slides (Ibidi) were coated with 2 μg/mL FN and incubated for 30 min 37 °C. Slides were blocked in 1 % BSA in 1X PBS for 30min at 37 °C. Cells were plated at 6 000 cells cm2 and incubated overnight. Before imaging, cell-permeable Hoechst (0.05 nM) (Thermo Scientific) and HEPES (10 mM) (Gibco) were added to the media.

### Immunostaining

Immunofluorescence staining was performed as described (38) with minor modifications. Briefly, cells on μ-slides were fixed and permeabilized with 4 % PFA, 0.5 % Triton X-100 and Alexa Flour 647 Phalloidin (Thermo Scientific) in cytoskeleton buffer (CT; 10 mM 2-(N-morpholino)ethanesulfonic acid, pH 6.1, 138 mM KCl, 3 mM MgCl, 2mM EGTA) for 15 min at 37 °C. Free aldehydes were reacted with 10 mM glycine for 15 min at room temperature (RT) and cells were washed with TBS at RT. Cells were incubated with primary antibodies (Mouse anti-paxillin; 1:350, BD biosciences) in a blocking solution (2 % BSA in TBS-T) supplemented with phalloidin (1:500) overnight and then washed and incubated with secondary antibodies (Alexa Flour 488 conjugated goat anti-mouse; 1:500, Invitrogen) in blocking solution and phalloidin (1:500) for 1h at RT. Cells were washed with TBS-T and blocked in blocking solution (2 % BSA in TBS-T) before imaging.

### Live fluorescence microscopy

Images were acquired using an inverted Nikon Ti2-E wide-field fluorescence microscope with a Nikon Plan Apo λ 10× 0.45 numerical aperture (NA) objective lens and Perfect Focus System (PFS) for maintenance of focus over time. Excitation and emission light were passed through DAPI (Exc. 379-405nm, Em. 414-480nm), FITC (Exc. 457-487nm, Em. 503-538nm), TRITC (Exc. 543-566nm, Em. 582-636nm) and Cy5 (Exc. 590-645nm, Em. 659-736nm) filter cubes, all from Semrock. Samples were kept in a humidified atmosphere at 37 °C and 5 % CO2 using an environmental chamber (Okolab). Images were acquired on a Nikon DS-Qi2 CMOS camera. The imaging of the samples was automated by generating stage positions covering the sample area using JOBS (NIS-Elements extension; Nikon) and a Nikon TI-S-ER motorized stage with an encoder. For time-lapse imaging, images were collected every 10 min for 100 min. The same system was also used with a Nikon CFI SR Plan Apo IR 60XAC WI / 1.27NA with a software-driven TI2-N-WID Water Immersion Dispenser for imaging of fixed samples or time-lapse imaging of live cell migration every 3 min for 3 hours.

### TIRF microscopy

TIRF microscopy was performed on an inverted TIRF microscope (Ti-E; Nikon) using a Nikon CFI Apo TIRF 100X Oil /1.49 NA. Excitation and emission light was passed through Continuous Storm (Nikon; Exc. 387-417nm and 557-570nm Em. 422-478nm and 581-625nm), FITC/TRITC ET (Exc. 450-490nm, Em. 500-530nm) filter cubes. Images were acquired on a Prime 95B 22mm sCMOS camera (Teledyne photometrics).

### N-SIM microscopy

SIM microscopy was performed on an inverted N-SIM microscope (Ti-E; Nikon) using a Nikon CFI SR APO TIRF 100X Oil / 1.49 NA excited by a Nikon LU-N3-SIM 488/561/640 laser unit. Excitation and emission light was passed through SIM 405/488/581/640 (Exc.484-496nm, 557-567nm and 629-645nm) filter cubes and N-SIM 488 BA (Em. 500-545nm), N-SIM 561 BA (Em. 570-640nm) and N-SIM 640 BA (Em.663-738nm) emission filters, all from Nikon. Images were acquired on an ORCA-Flash 4.0 sCMOS camera (Hamamatsu Photonics K.K) and the images were reconstructed with Nikon’s SIM software on NIS-Elements Ar (NIS-A 6D and N-SIM Analysis). Standard fluorescence microscopy was also performed on both the SIM and the TIRF systems with a Nikon CFI Plan Apochromat λ 10X / 0.45NA objective by generating stage-positions in the corners of the sample area using JOBS and a TI-S-ER motorized stage with encoder. For wide-field excitation, a SPECTRA X light engine ® (Lumencore inc) was used. All systems were controlled using NIS-Elements (v. 5.21.02 or later).

### DDM Framework

A lightweight framework was developed in Julia (v.1.6.0) and deployed on a local server to handle image analysis and data informatics. Image analysis packages with a shared schema were developed (see image analysis) and loaded into the framework. For each experiment, the corresponding analysis package was initiated with an experiment-specific configuration. Images were transferred to the server using an in-house developed command line interface (CLI; available on Github) for analysis. NIS Elements JOBS was used to query the server through the CLI for analysis output. See Supplementary Fig. 1 and Supplementary Note 1-4.

### Image analysis

All described image analyses were written in Julia (version 1.6.0) and are available on Github. See Supplementary Note 1.

### Transfection plugin

Low magnification (10X) images of multi-labeled fixed HeLa cells were acquired as described above. Image background segmentation was performed on the DAPI channel (cell nuclei). Using the foreground segments as seeds, a local thresholding (Otsu’s thresholding) window was used to generate segments of the cell fluorescence in each channel. Segments overlapping the seeds were selected and intensity features (i.e. mean, median, standard deviation) were measured and normalized to the mean sample background. Cells were sampled for data-dependent imaging based on the local relative intensity for every label. See Supplementary Note 5.

### Migration plugin

Low magnification time-lapse images of stable mKate-paxillin expressing H1299 cells were acquired as described above. Background segmentation was performed on the DAPI channel (cell nuclei). Features describing cell migration (e.g. mean speed, meandering index) were obtained by tracking the coordinates of the nuclei and finding the optimal assignment of coordinates, in terms of least total Euclidean distance, between each pair of frames using the Munkres algorithm (39). Background segmentation was performed on the TRITC channel (paxillin) to obtain intensity features (i.e. mean, median, standard deviation). Cells sampled from the top and bottom 10th quantiles in terms of mean speed were selected for data-dependent imaging. See Supplementary Note 5.

### Correlative imaging plugin

Low magnification time-lapse images of H1299 WT cells were acquired as described above (see live fluorescence microscopy). Prior to imaging the last frame of the time-lapse, the fixation and permeabilization solution (described under immunostaining) was added to samples to stop cell migration and record the last known relative stage coordinate of each cell. For the analysis: cell migration was analyzed as described above. On a secondary system (i.e. SIM or TIRF), JOBS was programmed to image randomly generated points in each corner of the sample in widefield (see TIRF/N-SIM microscopy). The cell nuclei were segmented for each image and the triangle between the coordinates of each nucleus and its 2 closest neighboring nuclei were calculated and normalized to a set of rotation and scale invariant measurements. Each set of measurements was then compared to the same measurements made on the last time point of the migration data. A RANSAC algorithm sampled the measurements based on the similarity between the two datasets (migration data and sampled image) and calculated an affine transformation matrix. If sufficient cells (>30 %) in the sampled image could be mapped onto the migration data, each matched cell pair was stored, and the process was repeated until 3 corners of the sample were matched. A final affine transformation matrix was calculated using matched pairs of all 3 corners (see Supplementary Fig. 5) and applied to the migration dataset. Cells were sampled as described above (see migration module) for TIRF and SIM imaging. Background segmentation was performed on the DAPI channel (cell nuclei). The foreground segments were then retouched manually to segment and separate the complete cells. The FITC channel (paxillin) was tophat transformed with a circular kernel with a radius of 6 pixels after which Otsu’s thresholding was performed to obtain morphometrics of the focal adhesions. Cell morphometrics (e.g. area) was obtained by background segmenting the TRITC channel (cell actin). See Supplementary Note 5.

### Automated data-dependent microscopy

Image features extracted from each experiment (e.g. cell morphometrics, fluorescence signal) were stored in a database and the recorded stage position of each data point was accessed through a query using the CLI. Each stage position was recorded to a local file on the microscope station and loaded into NIS-Elements for imaging using JOBS. The stage positions were imaged differently depending on the microscope: (a) for the live-fluorescence microscopy, the JOBS was programmed to switch to the Nikon CFI SR Plan Apo IR 60XAC WI / 1.27NA objective, apply water using the water dispenser, perform autofocus and subsequently image each point, (b) for the TIRF and SIM microscope, the JOBS was programmed to let the user switch to the respective Nikon CFI Apo TIRF 100X Oil / 1.49 NA objective, manually apply immersion oil and calibrate the sample to its original position, perform autofocus and image each point

## Statistical analysis

Statistical analyses were performed using Prism (GraphPad).

## Author contributions

Conceptualization: OA, JKA and PN. Experimentation and data analysis: OA and JKA. Writing original draft: OA and PN. All authors contributed to reading and editing the final manuscript.

## Acknowledgements

We thank the Lund University Bioimaging Centre (LBIC) for use of fluorescence microscopes. We thank the Emil and Eva Cornell Foundation, The Swedish Research Council (Vetenskapsrådet), The Swedish Cancer Foundation (Cancerfonden), The Strategic Research Foundation (SSF) and Per-Eric and Ulla Schyberg’s Foundation for funding.

