## Supplemental files for "Data-driven microscopy allows for automated targeted acquisition of relevant data with higher fidelity"

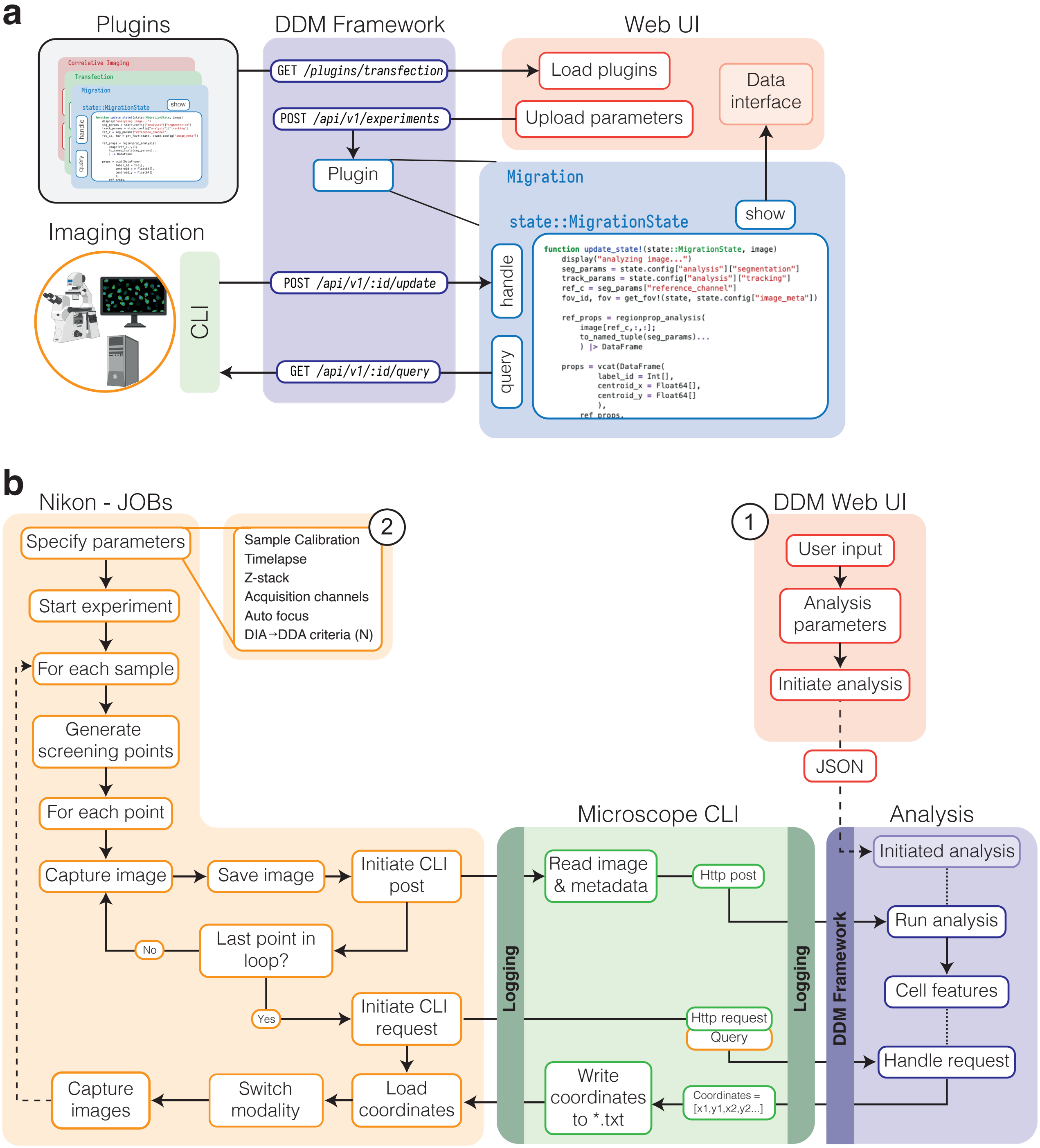


**Fig. 1. Data driven microscopy schematics. a.** The DDMFramework is light-weight, hosted on a local server, and responsible for the analysis of image data. Each analysis (plugin) is loaded into the framework by the user during the initiation step (step 1 in b) which can be interfaced though the framework. The analysis extends the framework with a handler (analysis), query (schema for accessing data) and show (user-data interaction). **b.** In more detail, the DDM-Framework hosts a simple UI through the browser where the user specifies parameters to be used during the analysis. Next, the user specifies how the microscope should perform the data-independent acquisition (DIA) and data-dependent acquisition (DDA). Each image acquired is stored locally and the CLI is triggered, posting the image to the DDM-Framework. When all images have been taken, a request is made using the CLI for coordinates from the framework. Received coordinates are stored in a `.txt` file and loaded into the microscopy-pipeline. Then, the DDA modality is initiated and images are acquired.

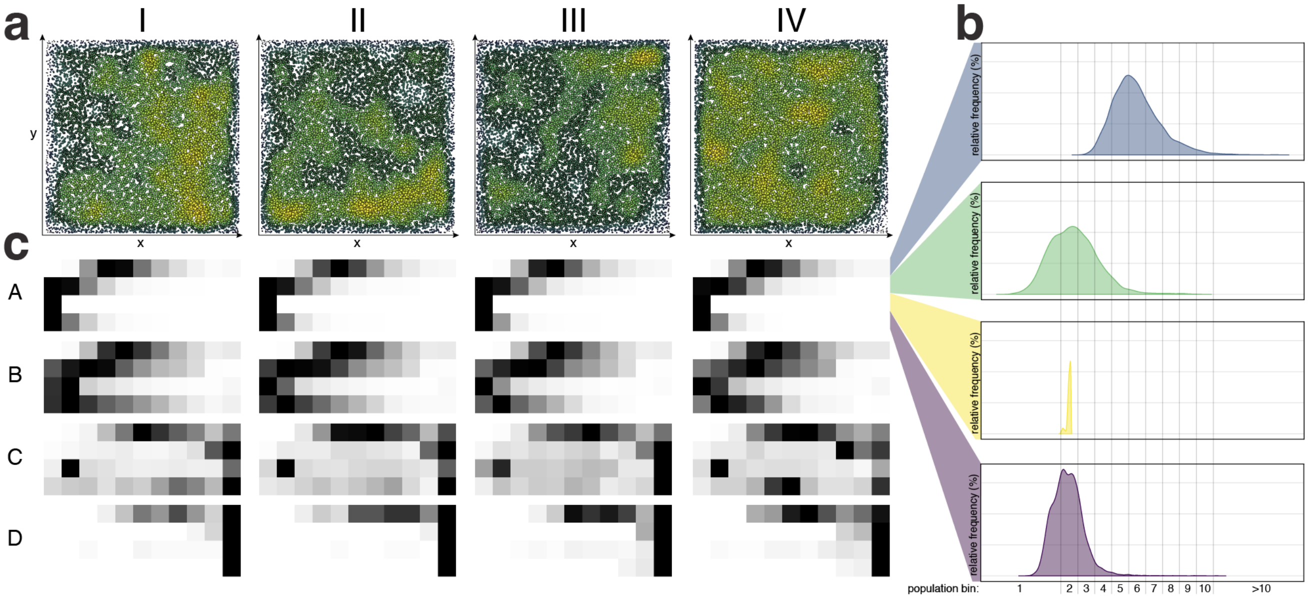


**Fig. 2. Phenoprints allow for compact visualization of phenotype characteristics. a.** Density plot of the spatial distribution of HeLa cells within four replicate samples. **b.** Histograms of the cell population distribution with regards to level (here normalized to relative bins) of four different fluorescent markers. **c.** Phenoprints were developed to visualize phenotype characteristics of a given population or subpopulation. Phenoprints consists of multiple feature vectors condensed into binned groups (panel b). Each bin is populated according to the population of cells within a given interval and feature. By stacking multiple feature vectors, it is possible to visualize the distribution of different population over selected features and use it for cross-sample comparison (panel c). The different phenotypes (A,B,C,D) shown above display small cross-replicate differences (horizontal comparison), but large differences across phenotypes (vertical comparison). Data is based on a subset of cellular phenotypes analyzed in Figure 3.

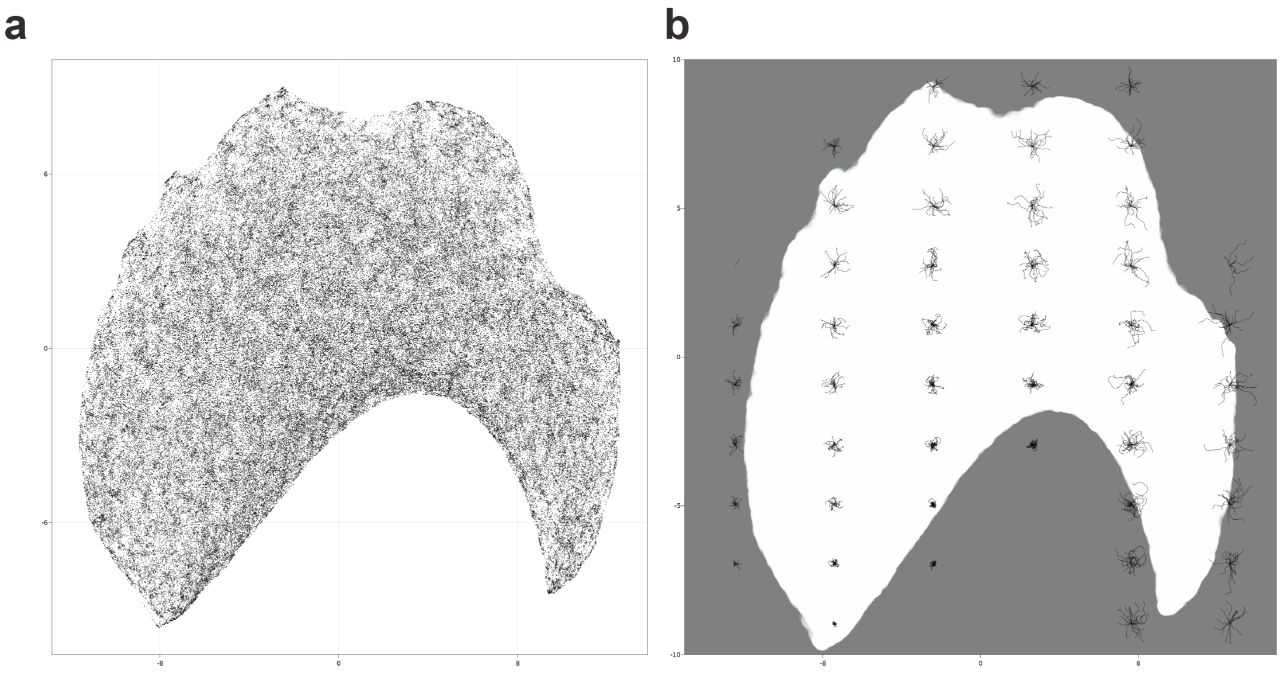


**Fig. 3. UMAP and cell migration tracks. a.** A UMAP coordinate space of migratory H1299 cells. Single cell data of migration speed (mean and standard deviation (SD)), migration persistence (angle; mean and SD) and total displacement over time (sum) was used to build the UMAP space. **b.** The UMAP space in a smoothed and overlayed with the migration tracks of cells binned $(6 \times 10)$ according to the coordinate space of the underlying UMAP.

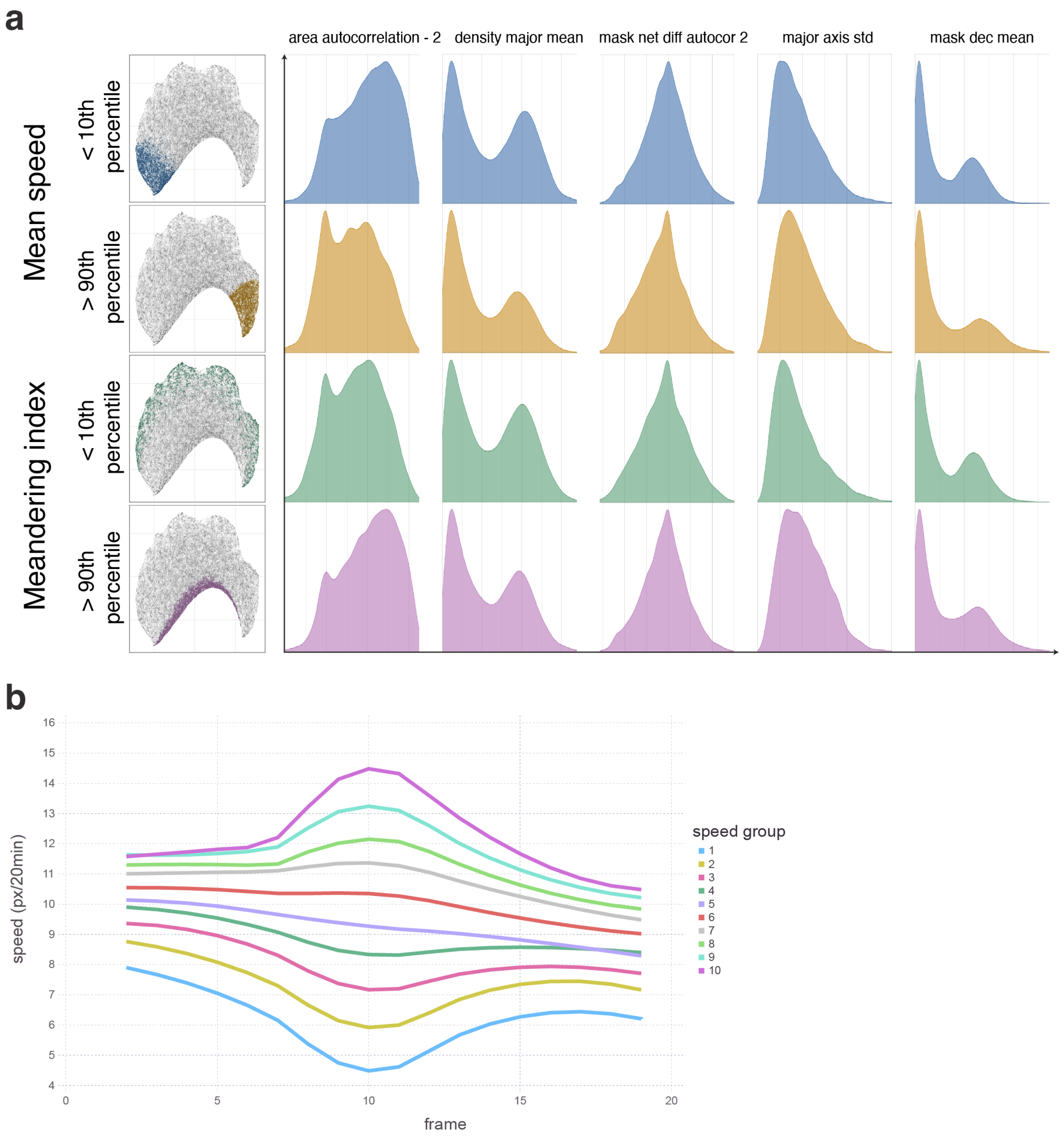


**Fig. 4. Single cell features of migratory H1299 cells. a.** Single cell feature distributions (area autocorrelation - 2, density major mean, mask net diff autocorrelation - 2, major axis std, mask dec mean) grouped according to top and bottom (>90th and <10th, respectively) quantile. Little differences was seen in distributions between the groups. Highlighted is the respective region of the groups in the underlying UMAP space (See Supplementary Figure 3). **b.** Grouping cells based on speed showed a inter-group consistency. Cells were grouped according to their quantile (10 total) in terms of speed in frame 10 (200min). All groups show a tendency to drift towards the population average in terms of speed. Slow cells (groups 1–5) tended to be faster and fast cells (groups 6–10) tended to be slower prior to grouping. The groups stayed within their speed hierarchy (e.g. fast cells remained faster than slow) after labeling with no overlapping inter-group with the exception of group 4 and 5.The decline in mean speed over time after grouping could be explained by overpopulation and the decline in free migration space.

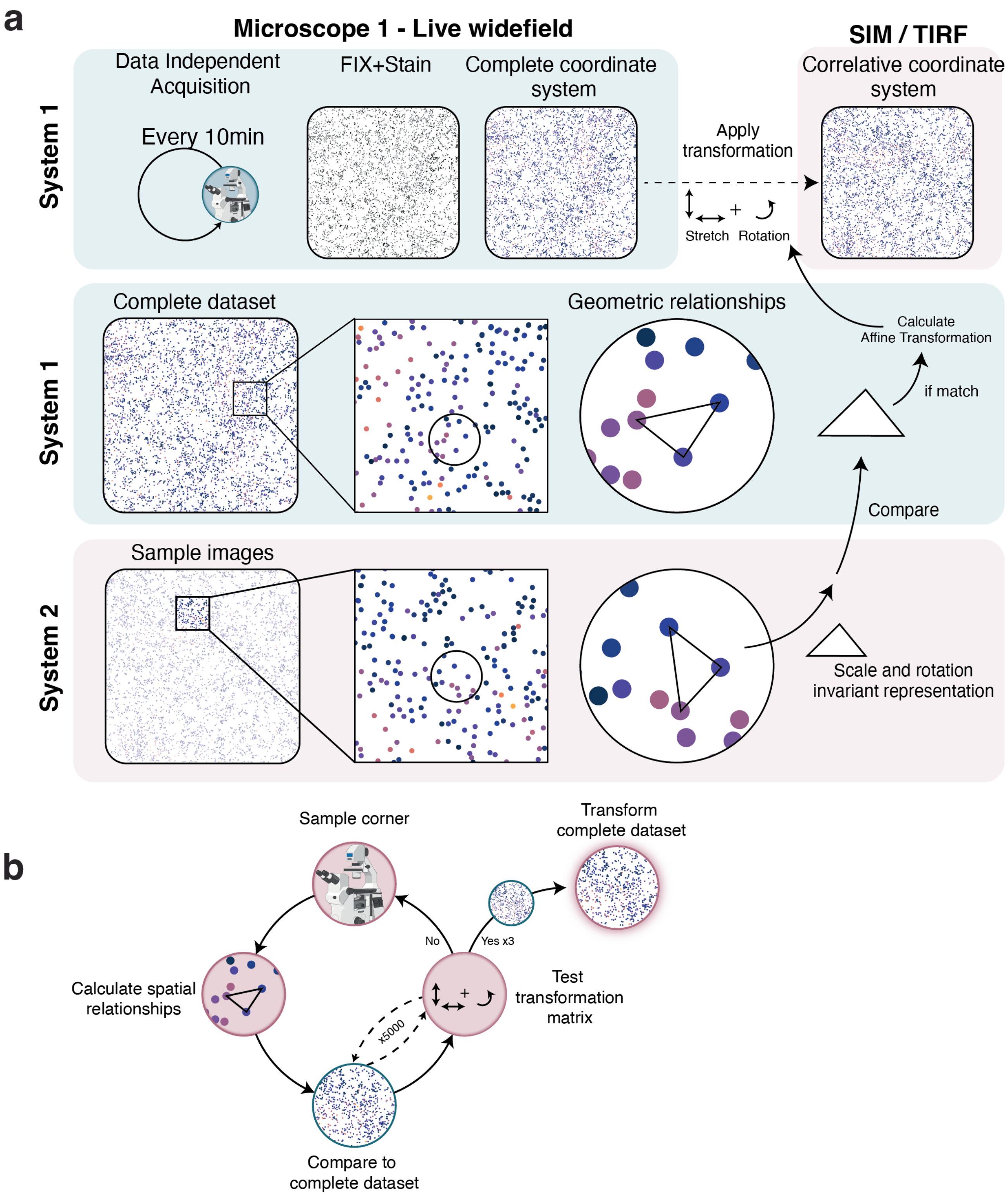


**Fig. 5. Correlative imaging procedure. a.** Migratory H1299 cells were imaged in DAPI (Hoehcst - Nuclei) and DIC every 10min for 6hours. To avoid post-acquisitional migration, cells were fixed and stained during the last timepoint, resulting in a complete dataset in system 1 with fixed coordinates in the alst timepoint. The triangle features (i.e. angles) were measured for each cells and its closest 3 neighbors. Through random acuisition of the same sample in system 2, the spatial features can be matched to system 1. If sufficient (>30%) cells match, the affine transformation is calculated for matched cells, which in turn can be applied to the complete dataset from system 1. **b.** In more detail, a random field of view is acquired for each corner of the sample. A RANSAC algorithm samples the coordinates based on the simularity of the scale and rotation invariant representation and generates a local affine transformation that best match the sampled field of view (system 2) to the complete dataset (system 1). If sufficient (>30%) coordinates (system 2) can be transformed and mapped onto the complete dataset (system 1), the matched coordinates are stored and a new corner is sampled. This procedure is repeated until 3 corners have been successfully calibrated to the complete dataset. Upon meeting this criteria, a final affine transformation is calculated using all the matched coordinates from all corners and applied to the complete dataset (system 1). This enables the sampling and targed acquisition of coordinates from system 1 in a second system.

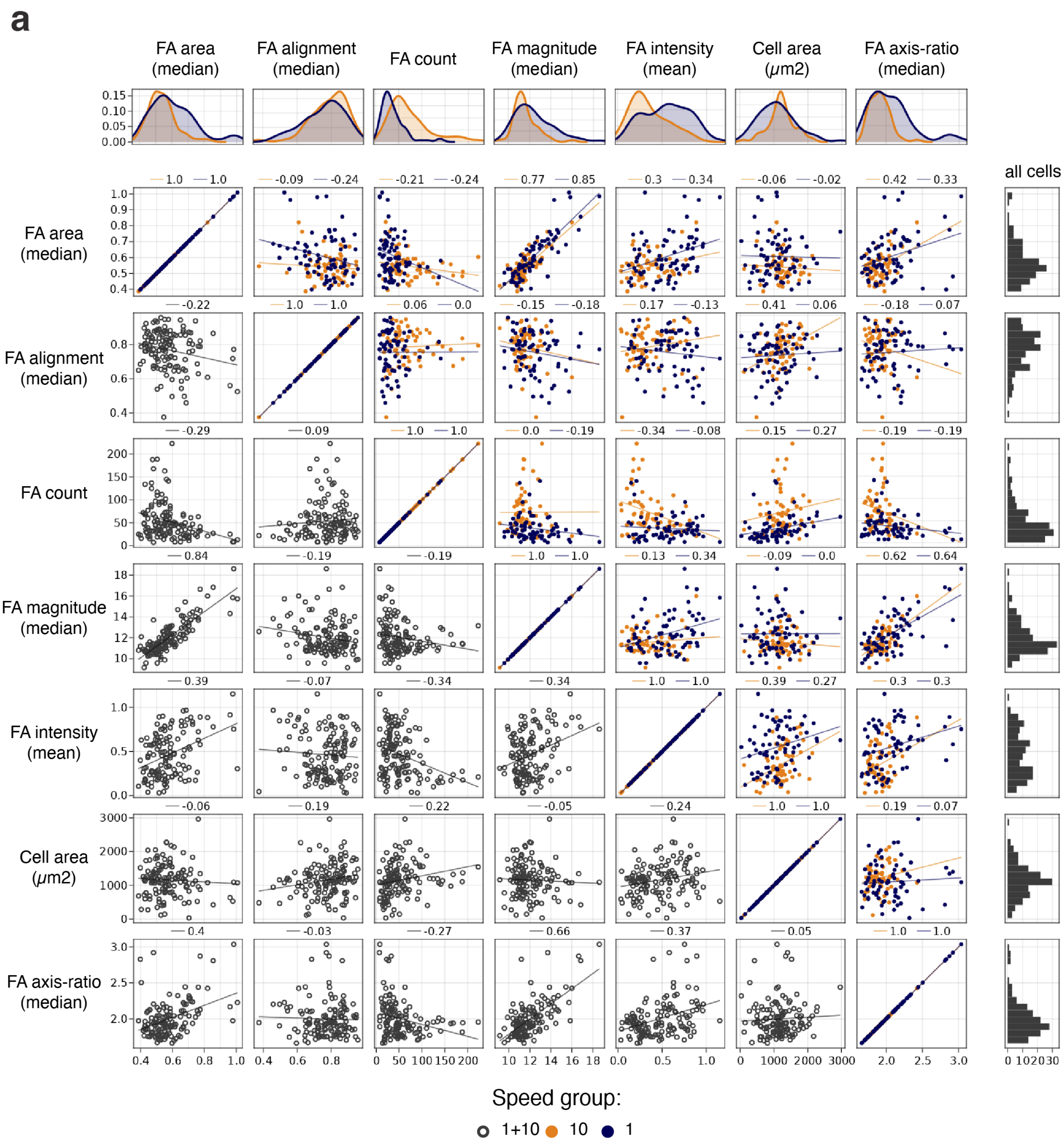


**Fig. 6. Visual exploration of correlations between focal adhesions and cell features. a.** Focal adhesion (FA; area, alignment, count, magnitude, intensity and axis ratio; timepoint=100min) and cell (area) properties from cells migrating in the top and bottom percentile(90th and 10th percentile, respectively; speed group 10 and 1) in terms of mean speed. There is some separation between the two groups in the different features (distribtion plots;top row). Correlations between the different features in the groups can be seen in the scatter plots (centre-plot; spearman correlation coefficient declared above each scatter plot).

### Supplementary Note 1: Github

The DDM framework can be accessed from: https://github.com/NordenfeltLab/DDMFramework

The individual plugins can be accessed from:

**DDMTransfection**
https://github.com/nordenfeltLab/DDMTransfection.jl

**DDMMigration**
https://github.com/nordenfeltLab/DDMMigration.jl

**DDMCorrelativeImaging**
https://github.com/nordenfeltLab/DDMCorrelativeImaging.jl

The command line interface responsible for microscope-framework communication can be accessed from:

https://github.com/NordenfeltLab/ddm_microscope_cli

### Supplementary Note 2: Hosting a DDM server

To host a DDMFramework, download the compiled version from github (see above) and run locally. To run plugins written in Python or Julia, simply store the corresponding python script or julia package in the *DDMFramework/plugins/* folder. Available plugins are loaded during initialization and are accessible on:

*localhost:4443/plugins*

Which opens a simple UI that displays the plugins. To initiate experiments on the plugin, parameters are posted in the form of JSON. See below for further information about each plugins parameters. Posting parameters is done through:

*localhost:4443/plugins/plugin_name/*

Each upload/post will create an experiment with increment id's starting with 0.

*localhost:4443/plugins/plugin_name/experiment_id/*

It is possible to have multiple experiments running with different parameters. Information about the running server and its plugins are available on the main page.

*localhost:4443/*

### Supplementary Note 3: Automation of microscopy.

Automation of the microscopes was performed using the latest version of NIS-elements AR and JOBs. Different JOBs programs were developed for each application (i.e. DDMTransfection, DDMMigration, DDMCorrelativeImaging). For interactive HTML versions of the JOBs programs, see Supplementary Files 1-3.

### Supplementary Note 4: Microscope-server communication

In order to communicate between hardware and the plugins running on DDMFramework, we developed a simple command line interface (CLI; written in Nim programming language, https://nim-lang.org/). The CLI is responsible for posting data, most commonly image-data, and querying data from the plugins. Every response from the framework is stored in a JSON text file. Additionally, the CLI logs all communication to a `.txt` file. For further information about the CLI, we refer to the github page above.

For this paper, DDM was performed on hardware controlled through Nikon's NIS-elements. Executing tasks with the CLI in the case of Nikon was handled through the internal macro language and JOBs (See Supplementary File 1-3).

**POST: send.** To send data, for instance image-data, to the framework, the CLI is called with the `send` command. The `send` command reads the acquired data in a directory and performs a HTTP POST to the specified address. `send` takes 5 arguments, being:

```
Usage:
  send [optional-params]
Options:
  -a=, --address=       "http://localhost:4443"  Server address, including
                                                 domain.
  -d=, --data_path=     "images"                 Filename of the data to be sent in root_dir.
  -r=, --response-path=  "reponse.txt"           Path to write response in root dir.
  -l=, --logging-path=  "log.txt"                Path to log file in root dir.
  --root_dir=           "./"                     Path to root dir, defaults to internal os.getCurrentDir().
```

The response from the framework is parsed as JSON and written to the declared response-file.

**REQUEST: queryDDM.** To request data from the running plugin on the framework, the CLI is called with the `queryDDM` command. In addition the the arguments of `send`, `queryDDM` takes 2 arguments:

```
  -q=, --queryString=    ""                      Query as argument.
  --queryPath=           ""                      Path to query as txt file (overrides queryString).
```

Received data is parsed as JSON and written to the declared response-file.

**Query Structure.** A simple custom query-language was developed to query data from the plugins. In the future, we intend to support multiple ways of querying data, including GraphQL (`https://graphql.org/`). Plugins can also opt for completely custom query-logic (see DDM Plugins, a short description).

The following query-structured is parsed:

$$plugin(filter, order)(columns\ldots)$$

Where:

***plugin*** Defines name of the DDM plugin to query.

***filter*** Defines filtering the data.

***order*** Defines the order of the data.

***(columns. . . )*** Defines returned columns of the data.

And the following operands have the structure:

***filter*** filter:{column:[op:operation, arg:[arguments]]}

***order*** order:[[type,column]]

Finally, (columns) specified the columns in the plugin to be returned. Each plugin declares what columns are accessible from queries.

For instance:

*...{migration(filter:{mean_speed:[{op:"gt",args:[12]}]},order:[["desc", "area"]])(stage_pos)}*

Which would yield a response from plugin *migration* containing all entries filtered on column *mean_speed* greater than 12 and descending order on area, as declared in the plugin *migration*. Multiple query operations (i.e. filter, order) can be performed on multiple columns in succession. For instance:

*filter:{mean_speed:[{op:"gt",args:[12]}],circularity:[{op:"ls",args:[0.6]}]}*

In addition, the query-parser supports sampling of the data. This can come in the form of limiting the data to a certain N entries in order, or sample N entries randomly from the dataset. This is declared before the other operands, but is computationally executed last. Extending the above query to sample 25 entries:

*migration(sample:25,filter...*

Or limit the dataset to 25 entries (in order):

*migration(limit:25,filter...*

The full path to the plugin-data, which the CLI directs the received query to, is the following:

*domain:port/api/v1/experiments/id/?query={plugin(filter,order)(columns...)}*

Where:

***domain*** Domain address.

***port*** Server port.

***id*** Experiment id.

***plugin*** DDM Plugin.

### Supplementary Note 5: DDM plugins, a short description

The DDMFramework package provides a general framework for data management and communication, allowing user-provided plugins to focus on image analysis and data processing. A plugin manages updating and querying data within the framework. The simplest way to create a plugin is by generating a dynamic plugin using the `multipoint` function.

```
multipoint(analyisis, name::String, keyfun; keytest=(==))
```

The first argument, `analysis`, is a function that receives acquired data and the plugin configuration. The `keyfun` argument is function used to generate a key from the data, typically from the metadata, sent by the client. This key is used to separate the data into bins, for instance images from the same field-of-view. To test if newly received data belongs to a bin, the function `keytest`, which defaults to equality (==), is used. The `name` is used when registering the plugin with the framework and rendering results. See below for an example of how a plugin can be created using `multipoint`, or on github (`https://github.com/NordenfeltLab/DDMFramework.jl`). Advanced users can get more flexibility by defining a type subtyping `AbstractPlugin` and fulfilling the plugin interface by extending `handle_update` and `query_state`.

```
handle_update(state::SimpleState, data::Dict{String, Any}) # Returns response
                                                          # and updated state
query_state(state::SimpleState, query) # Returns a String
add_plugin("simpleanalysis", SimpleState) # Loads plugin into the framework
```

**Developing DDM plugins using Python.** The DDMFramework package aims to support plugin development in multiple programming languages used in life-science. So far we provide basic support for Python.

**A. DDMTransfection.** DDMTransfection is a plugin that is used to estimate transfection of fixed cells in high-content screening. A background estimation algorithm (see github) is used to measure the background signal in each channel and segment the nuclei. The segmented nuclei acts as seeds for applying a window on each channel. In each winow, we perform Otsu's Thresholding to create segments of positive signal. If no segment can be made overlapping the seeded nuclei, we default to measure the intensity in the respective channel overlapping the nuclei-segment. Finally, each cell's intensity in each channel is normalized according to the sample background. To initiate the DDMTransfection plugin, a JSON-formatted text containing the following parameters is used:

```
{
    "channel_definitions" : [
        {
            "alias" : "Name of channel : String",
            "index" : "Index of channel : Int"
        }
    ],
    "comment" : "Repeat above object for each channel in the images"

}
```

**B. DDMMigration.** DDMMigration is a simple plugin used to segment and track nuclei in images over multiple fields of views and frames. A background estimation method is used to perform background segmentation of the nuclei. For tracking, the algorithm (1) was implemented in julia (for further reading, see DDMMigration on github). To initate the DDMMigration plugin, a JSON-formatted text containing the following parameters is used:

```
{
  "analysis": {
    "segmentation": {
      "function" : {
          "method": "Segmentation method*",
          "minsize": "Minimum size of objects (pixels)* : Int",
          "maxsize": "Maximum size of objects (pixels)* : Int",
          "..." : "Further analysis-specific parameters"
      },
      "primary_channel": "Index of primary image to segment : Int",
    },
      "tracking": {
          "maxdist": "Maximum distance-allowance for tracking objects : Float"
      }
  },
  "system": {
    "camera_M": {
      "a22": " : Float",
      "a21": " : Float",
      "a12": " : Float",
      "a11": " : Float"
    },
  }
}
```

*overrides default

**C. DDMCorrelativeImaging.** DDMCorrelativeImaging is an intermediary plugin that correlates points to that of another plugin. A scale and rotation invariant representation of each position (centroid) of an object in an image is created with the two closest objects. In the control dataset from the secondary plugin, the same methodology is applied to all objects. A RANSAC algorithm samples points from the control-dataset and the hungarian-algorithm for the linear assignment problem (MUNKRES) is performed for all points. The affine transformation from the best matched representations is tested on the sampled field of view (fov). If >30 % of the objects in the fov can be mapped onto the control-dataset, the matched representations are stored and the process is repeated until three sets of matched representations (>30 %) are found. A final transformation is calculated with from an aggregate of all the matched representations, and mapped onto the control-dataset. See Supplementary Figure 5 for more info. Finally, the DDMCorrelativeImaging plugin acts as an interface to the secondary plugin by parsing the queries. To initiate the DDMCorrelativeImaging, a JSON-formatted text containing the following parameters is used:

```
{
    "analysis" : {
        "plugin": "Name of plugin : String",
        "exp_id": "Experiment id : Int",
        "port": "Port to secondary plugin : Int"
    },
    "segmentation": {
        "function": {
            "method": "Segmentation method* : String",
            "minsize": "Minimum size of objects (pixels)* : Int",
            "maxsize": "Maximum size of objects (pixels)* : Int",
            "..." : "Further method-specific parameters"
        },
        "primary_channel": "Index of primary image to segment : Int"
    },
    "system": {
        "camera_M": {
            "a11": " : Float",
            "a12": " : Float",
            "a21": " : Float",
            "a22": " : Float"
        }
    }
}
```

*overrides default

### Example plugin in julia: NucleusProperties.jl

```julia
using Images
using DDMFramework
using RegionProps
using DataFrames
using Chain

otsu_segment(img) = img .> otsu_threshold(img)

function filter_objects_in_image(labeled_image, minsize, maxsize)
    sparse_lb = sparse(labeled_image)
    counts = countmap(nonzeros(sparse_lb))
    for (i, j, v) in zip(findnz(sparse_lb)...)
        if counts[v] < minsize || counts[v] > maxsize
            lb[i, j] = 0
        end
    end
    dropzeros!(labeled_image)
end

function simple_segmentation(img; minsize=150, maxsize=2000)
    labeled_image = @chain img begin
        otsu_segment
        label_components
        filter_objects_in_image(_, minsize, maxsize)
    end
    return labeled_image
end

multipoint("NucleusProperties") do image, config
    # Segmentation parameters
    seg_params = config["segmentation"]

    # Segment and filter objects on size in image
    labeled_image = simple_segmentation(image, to_named_tuple(seg_params)...)

    # Extract stats about our objects
    return regionprops(
        image,
        labeled_image;
        selected=unique(nonzeros(labeled_image)),
    )
end
```

### Example plugin in python: NucleusProperties.py

```python
import numpy as np
import pandas as pd
from skimage.filters import threshold_otsu
from skimage.measure import label, regionprops_table
from math import isclose

def filter_objects_in_image(labeled_image, minsize, maxsize):
    values, counts = np.unique(labeled_image, return_counts=True)
    count_dict = dict(zip(values, counts))

    def object_filter(lb):
        if lb > 0:
            c = count_dict[lb]
            if minsize <= c <= maxsize:
                return c
        return 0

    return np.vectorize(object_filter)(labeled_image)

def simple_segmentation(img, minsize=150, maxsize=2000):
    binary = otsu_segment(img)
    labels = label(img)
    return filter_objects_in_image(labels)

def keyfun(data):
    x = data["image"].Pixels[:Plane][1][:PositionX]
    y = data["image"].Pixels[:Plane][1][:PositionY]
    return (x,y)

def keytest(kleft, kright):
    atol = 1.2
    return all(map(isclose, kleft, kright))

def analyze(image, config):
    # Segmentation parameters
    seg_params = config["segmentation"]

    # Segment and filter objects on size in image
    labeled_image = simple_segmentation(image, **seg_params)

    # Extract stats about our objects
    return pd.DataFrame(regionprops_table(image, labeled_image))

add_plugin(multipoint(analyze, "NucleusProperties", keyfun))
```

### Online Supplementary File Notes

**Supplementary File Note 1 - DDMTransfection.** An interactive HTML version of the JOBS setup used for automation of microscopy control with the DDMTransfection module can be found at https://nordenfeltlab.github.io/DDMTransfection.jl/dev/microscopes/nikon/.

The acquisition on the microscope was automated using JOBS. Settings (e.g. capture definitions, autofocus settings, storage locations and run variables) for the acquisition are defined at the start of the program. The program performs the following for each well; the data-independent acquisition (DIA; left panel) generates points covering approximatly 85is performed on the first point. For each point, images are taken, stored locally and posted to the server through the in-house developed command-line-interface (CLI). After all points have been imaged, the CLI is initiated with a query through a macro. The response dictates if available coordinates are loaded into the program. The secondary imaging modality (60X water immersion objective) is initiated and autofocus and imaging is performed on each coordinate. Once all images are taken, the primary modality is initiated and the process repeats for the next well. DIA and DDA are performed in the same JOBS program.

**Supplementary File Note 2 - DDMMigration.** An interactive HTML version of the JOBS setup used for automation of microscopy control with the DDMMigration module can be found at `https://nordenfeltlab.github.io/DDMMigration.jl/dev/microscopes/nikon/`.

The acquisition on the microscope was automated using JOBs. Settings (e.g. capture definitions, autofocus settings, storage locations and run variables) for the acquisition are defined at the start of the program. The program performs the following for each well; the data-independent acquisition performs a timelapse and generates points covering approximately 45Autofocus and perfect-focus system is perform for the first point in the first timepoint to keep focus throughout the DIA. Images are captured for each point and saved to the defined storage location. A macro initiates the in-house developed command-line-interface (CLI) that posts the image to the server. After the timelapse is complete, data-dependent acquisition starts. Two macros containing two seperate queries are initiated, in this case for slow and fast migrating cells. For each query, if the response contains coordinates, they are imported. Autofocus is performed on each coordinate and saved. Lastly, images are taken for each coordinate in higher magnification for the duration of the timelapse. Once complete, the process is repeated for next well.

**Supplementary File Note 3 - DDMCorrelativeImaging.** Interactive HTML versions of the JOBS setups used for automation of microscopy control with the DDMCorrelativeImaging module can be found at `https://nordenfeltlab.github.io/DDMCorrelativeImaging.jl/dev/microscopes/nikon/`.

JOBS was setup to automate the acquisition and calibration of the system. Settings are defined at the start of the program. Data-independent acquisition was performed on system 1. All wells were imaged every 10min for 100min total. Resulting data was uploaded to the server and analyzed. On the other microscopy systems, the data-dependent acquisition was performed. For each well, a point-set of the corners are generated. For each corner, random points are sampled and imaged, posted to the server through a in-house developed command-line-interface (CLI) using a macro. Subsequently, a request is made through the CLI with another macro. If the response contains '1' without coordinates, we break from the random points and continue to the next corner. This is repeated until the response contains coordinates. Once complete, the user is prompted to switch to the acquisition objective and re-align the sample. Finally, each point is imaged with auto-focus in high magnification.
